# Dissecting mycorrhizal fungal trait variation, its genetic basis and trade-offs

**DOI:** 10.1101/2025.10.08.681103

**Authors:** Erica McGale, Jamille Viray, Philip Gwyther, Ian R. Sanders

## Abstract

Despite the importance of arbuscular mycorrhizal fungi (AMF) in worldwide ecosystems, there is relatively little controlled phenotyping of their intraspecific fungal trait diversity, which limits our determination of how AMF traits link to their genetic differences. Here, we monitor 33 *in vitro* fungal traits in a population of 48 genetically diverse isolates of *Rhizophagus irregularis*, over 18 weeks, across plate compartments with and without direct plant root influence. This study reveals that intraspecific AMF variability is extensive, and that *R. irregularis* traits are mostly positively correlated, especially in root-free environments. The few trait trade-offs (negative correlations) found existed in proximity to the host root. Given the large screening population and the variance decomposition method of repeatability, we find that the genetic individuality of isolates significantly explained variation in approximately half of the AMF traits measured, and especially those related to hyphal growth, branching structures and reproduction near plant roots. Especially, differences in the frequency and fineness of branching right after growth from the plant host, which did not correlate to nearly any other traits, were explained by isolate identity (*i.e.*, genotype). The new information we reveal on *R. irregularis* trait relationships, over time and in different environments, as well as trait links to genetic differences among isolates, provides the foundation for future understanding of AMF functional diversity. Our genetic findings also provide critical groundwork for genome wide association studies on the non-plant-related traits of *R. irregularis*.

## INTRODUCTION

Arbuscular mycorrhizal fungi (AMF) are endosymbionts found in the soils of every continent [1] and are known to interact with a majority of plant species [2]. Over the last fifty years, the positive influence of AMF on the resilience and productivity of ecosystems has been increasingly supported [3,4]. AMF are mutualists of plants, establishing in the root cortex, and extending from there to increase the soil volume from which their plant partners can access nutrients, water and microorganisms [5,6]. The AMF gain energy and nutrients from their hosts to use for their own growth and reproduction [7,8]. Beyond their base dependence on plants, AMF growth traits are predicted to be influenced by many factors, from plant to fungal genetics, and environmental conditions [9,10]. However, few studies present direct evidence on how these variables alter AMF growth strategies during symbiosis [11].

Considerable advances have been made in establishing the molecular mechanisms of the initial interaction between a plant and AMF partner [8], as well as their resource exchanges [9,10]. In comparison, much less is known about genetic differences in AMF life history traits. Primarily, this is due to the fact that most AMF fungal traits are compared among species [11,12]. This makes the identification of shared genetic contributors across isolates hard to establish, which in turn makes it difficult to robustly predict the effects of AMF inocula. In agricultural settings, studies have reported negative, negligible and positive yield changes produced after inocula of various AMF species are added to soils already containing native AMF [13–16]. It is known that inocula of the same AMF species, containing different isolates, can cause very large differences in field crop yields [17–19]. The variation in plant yields afforded by different AMF isolates of a same species, and known higher within species functional variability, suggests that the relevant level to study AMF traits during symbiosis may be intraspecifically rather than interspecifically [20–22].

Recent papers reviewing trait-based frameworks for AMF highlight the scarcity of AMF trait characterizations that do not consider these fungi mainly for their value in relation to plants [11,23]. A better understanding of the contributions of fungal genetics to AMF growth traits, independent of benefits for the plant, is necessary, as fungal traits only partially correlate with effects of plant diversity [24]. This would be best achieved with the study of a population of a single species of AMF (as in [25] but extending from plant-related fungal traits). *Rhizophagus irregularis*, the model AMF species, is ideal for studying the drivers of intraspecific, during-symbiotic fungal traits in a controlled environment. Many *in vitro* cultured isolates of this AMF exist, several with previous characterizations at the genetic and epigenetic levels [26–31]. This species is also relevant as it is commonly found in many terrestrial ecosystems [32], is a popular component of commercial AMF inoculums [33] and is already cultured in a well-developed *in vitro* system in which fungal trait explorations have been shown to be possible [34–36].

In previous work, Savary and colleagues [34] found significant differences in the extraradical hyphal density of *in vitro R. irregularis* isolates according to their classification into phylogenetically distinct clades. Kokkoris, Miles and Hart [35] showed significant differences among early growth and reproduction traits (up to 35 days) of 11 *in vitro R. irregularis* isolates. With 10 of these 11 isolates, Serghi *et al.* [36] studied germination (asymbiotically, up to 30 days) in addition to some hyphal exploration traits (during symbiosis, up to 30 days after crossing into a hyphae-only petri dish compartment) and characterized the influence of plant host and fungal nuclear organization (homokaryotic or dikaryotic) on the growth of these *R. irregularis* isolates. Most recently, Oyarte Galvez and colleagues developed a high throughput system for tracking half a million fungal growth nodes in petri dishes and characterized network-like structures and cytoplasm fluxes in two *R. irregularis* and one *R. aggregatum* isolates [37].

To date, no studies have used a large enough isolate population to characterize AMF traits significantly influenced by isolate identity (i.e., their genetic individuality) versus other fungal, temporal, or environmental characteristics (e.g., isolate age, timing of growth or interactions with space). A large, diverse number of studied isolates could also allow the novel determination of correlations among traits measured within a single species. As of yet, most studies of potential AMF trait trade-offs are only reported at the species level, within the competitor-stress tolerator-ruderal (C-S-R) paradigm [38]. Uncovering the variation, correlations among and genetic basis of *R. irregularis* fungal traits during symbiosis could inform our understanding of the adaptation potential of a single species of AMF. Additionally, understanding which AMF traits are explained by genetic differences, and to what extent, is a prerequisite for genome wide association studies (GWAS) on those traits, leading to discovery of genes and regions of the AMF genome involved in fungal processes during symbiosis.

Here, we conducted an *in vitro* screening of traits in 48 isolates of *R. irregularis*, including phenotyping in compartments with and without plant material. We recorded 33 AMF traits related to hyphal and spore appearance and expansion. We also measured one plant trait; the plant material was otherwise controlled. We followed the isolates for 18 weeks and included a spatial and timed trial to detect minute influences on isolate traits. We included two outgroup *R. proliferus* isolates for comparison. We report new insights into the extent of variation and correlation of fungal traits in a single AMF species. We use a variance decomposition analysis (e.g., repeatability, [39]) to conclusively dissect the contribution of isolate genetic individuality, versus other attributes, on *R. irregularis* fungal traits during symbiosis. Our work thus provides the first understanding of how and when genetic determinants dominate AMF growth and justifies future work like GWAS.

## MATERIALS AND METHODS

### Fungal material and growth conditions

All isolates used (48 *Rhizophagus irregularis*, 2 *R. proliferus* serving as an outgroup) originated from *in vitro* cultures maintained at the University of Lausanne ([34]; Tab. S1). For the main trial, they were cultivated using a standard, sterile two-compartment petri dish system ([40]; Fig. 1A, n = 4 per isolate). One week after plant plating, a fungal plug containing ∼100 spores was applied on the carrot plug. The age since the last subculture and the general hyphal density of the fungal plug were noted to include as potential random factors (Tab. S2). Plates were sealed with parafilm and placed upside down in incubators at 25°C, without light, for the entirety of the trial.

**Figure 1.**
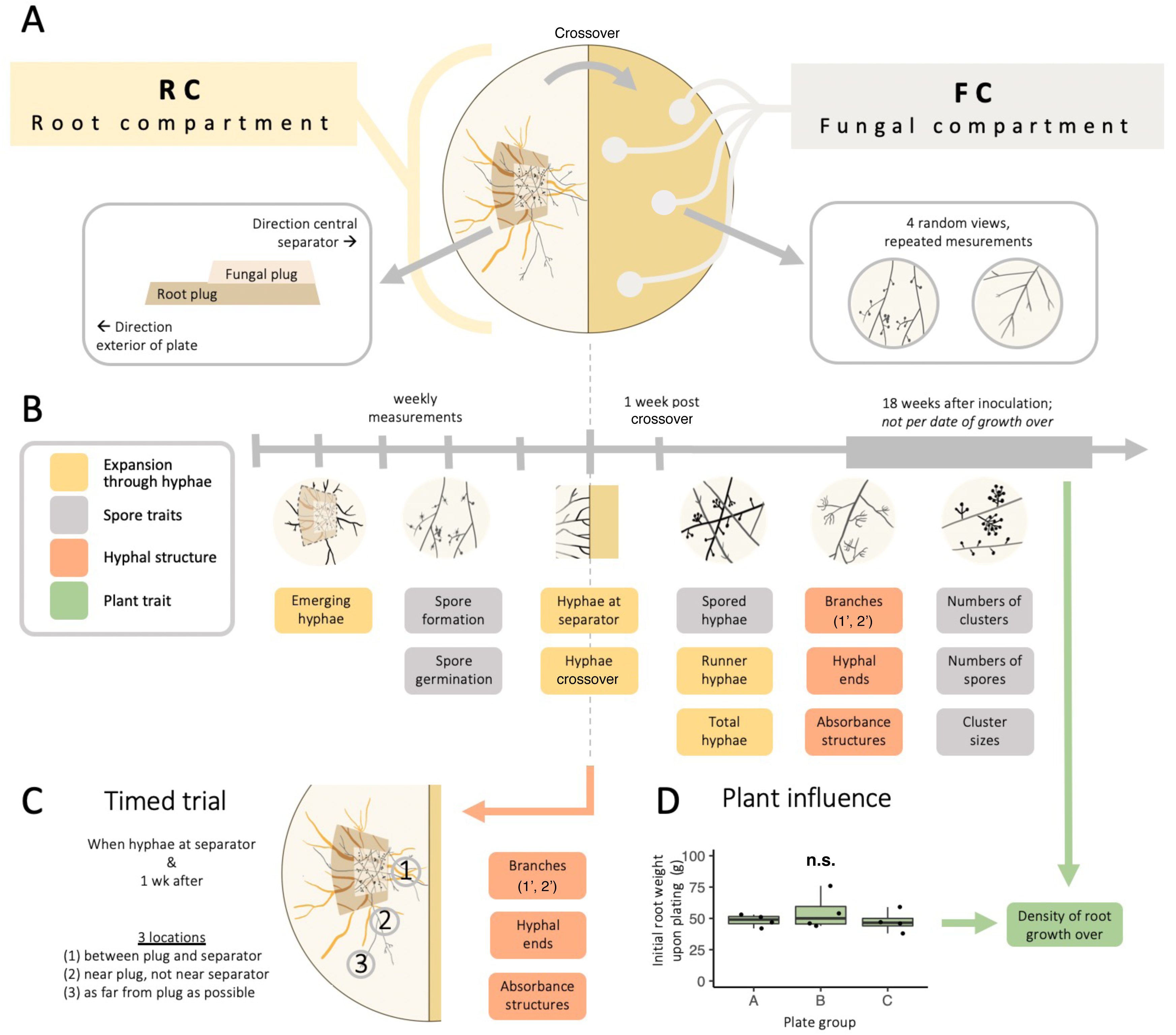
(A) Schematic of the experimental setup. The Root Compartment (**RC**, left) contains the starting fungal and root material, from which fungal hyphae (dark gray) and roots (orange) grow. The Fungal Compartment (**FC**, right) has a membrane (dark yellow) placed over the media to reduce carrot root growth but allow fungal hyphal growth into the media of the FC compartment. **(B)** Timeline and visualization of the measurements taken during the study. In the RC, weekly observations were taken in the whole compartment. In the FC, the traits were totaled from four random views within the compartment (**A**, right) at two time periods: 1 week after the first hypha of each plate grew over the central separator (**A**, “Crossover”), as well as during a single final measurement period at 18 weeks post inoculation. **(C)** Schematic of a timed trial on hyphal traits in the RC conducted when the first hypha touched the central separator. **(D)** Weight (g ± SE) of the initial amount of plant material used to inoculate. **n.s.** indicates not significant.

To explore the replicability of the experiment, a second trial with identical plating conditions was conducted with ten phenotypically diverse isolates (Tab. S1): nine isolates from the forty-eight of the first trial, and one additional isolate named *SAMP7*. As round number did not explain most trait variances for the isolates present in both trials (Tab. S3), data from both rounds were combined for all further analyses.

### Trait measurements in the root compartment (RC)

Over 16 weeks, weekly stereomicroscopic observations were made in the root compartment (RC, Fig. 1B) for numbers of hyphae emerging from the plugs, numbers reaching the middle separator of the two-compartment petri dish, and of new spores and their germinations (MZ125, Leica, Germany; 1.0x objective, 10x/21 eyepiece). Contaminated plates were removed, but their earlier measurements were kept. The number of active plates per isolate at key phenotyping timepoints are indicated in Tab. S4.

The first day of spore germination was recorded, as well as the number of days until hyphal or spore number reached a predefined threshold (50 for hyphae measurements, 250 for spores). All RC measurements were concluded one week after hyphae were first observed in the fungal compartment (FC). All 1^st^ day traits were directionally standardized to ensure low values can be interpreted as slower growth, and vice versa (*i.e.*, for 1^st^ day of spores germinating, a lower 1^st^ day originally indicated faster growth; we standardized this by subtracting the values from a common number, producing larger values for earlier, faster actions and smaller values for later, slower ones, without otherwise changing the data). The 1^st^ day measurements of hyphae reaching the separator and of germination were additionally normalized to the 1^st^ day of hyphae and spore emergence, respectively (by subtraction of the day of emergence). Once hyphae began gathering at the separator, the FC was checked for potential crossover (1^st^ day of Crossover).

A two-week timed trial assessed variability in RC hyphal structure traits. Counts of hyphal branches (primary, secondary), hyphal ends, and highly branched absorbance structures were taken at 20x from three RC locations with varying spatial constraints (Fig. 1C). The timed trial began for each respective plate when the first hypha reached the middle separator. The three different locations did explain RC hyphal trait variance (Tab. S5), but to be consistent with the treatment of FC hyphal measurements (where location was not a significant contributor, Tab. S6), timed trial trait values used for the main analyses were totals of each trait at the three RC locations.

### Trait measurements in the fungal compartment (FC)

FC growth was characterized in four random 40x views (2 near the plate edge, 2 near the center of the FC) to test edge effects as a random factor. View location did not explain trait variability (Tab. S6); the four views were thus combined for further analyses.

FC phenotyping was one week after each plate’s Crossover, including total numbers of hyphae (with and without spores), hyphal branching points, ends, and absorbance structures (Fig. 1B). Spore data included the number of spore clusters along the longest visible hypha, cluster size distributions, and total spores in the densest view area (80x magnification). A second FC measurement was taken when all plates had Crossover, at 18 wpi (i.e., not adjusted to Crossover date). The same views and traits were measured.

Relative carrot root growth into the FC was the only plant trait measured, at 18 wpi. Carrot material was otherwise standardized at the experiment start: the plant plug contained two thick roots (∼2 cm in length, shown to produce a consistent carrot tissue weight across plates: Fig. 1D).

### Data processing

All analyses were conducted in R version 4.0.5. Raw data and calculated AUCs (areas under the curve), after directional and emergence timing normalizations, were used for correlations (Figs. 2, 4, S3, S4), rankings (Figs. 3, S2), and non-metric multidimensional scaling, NMDS, analyses (Fig. S1). Non-gaussian trait and AUC data were log-transformed for repeatability analyses (Fig. 5; Tab. S3, S5, S6) to fit the assumption of data normality [41]. AUCs were generated from all the data points of select traits, per plate, taken over time after fitting each trait’s dataset to the best fit among a log, linear, quadratic or exponential model, and integrating this over 60 days (∼10 weeks) from the first occurrence of each trait.

**Figure 2.**
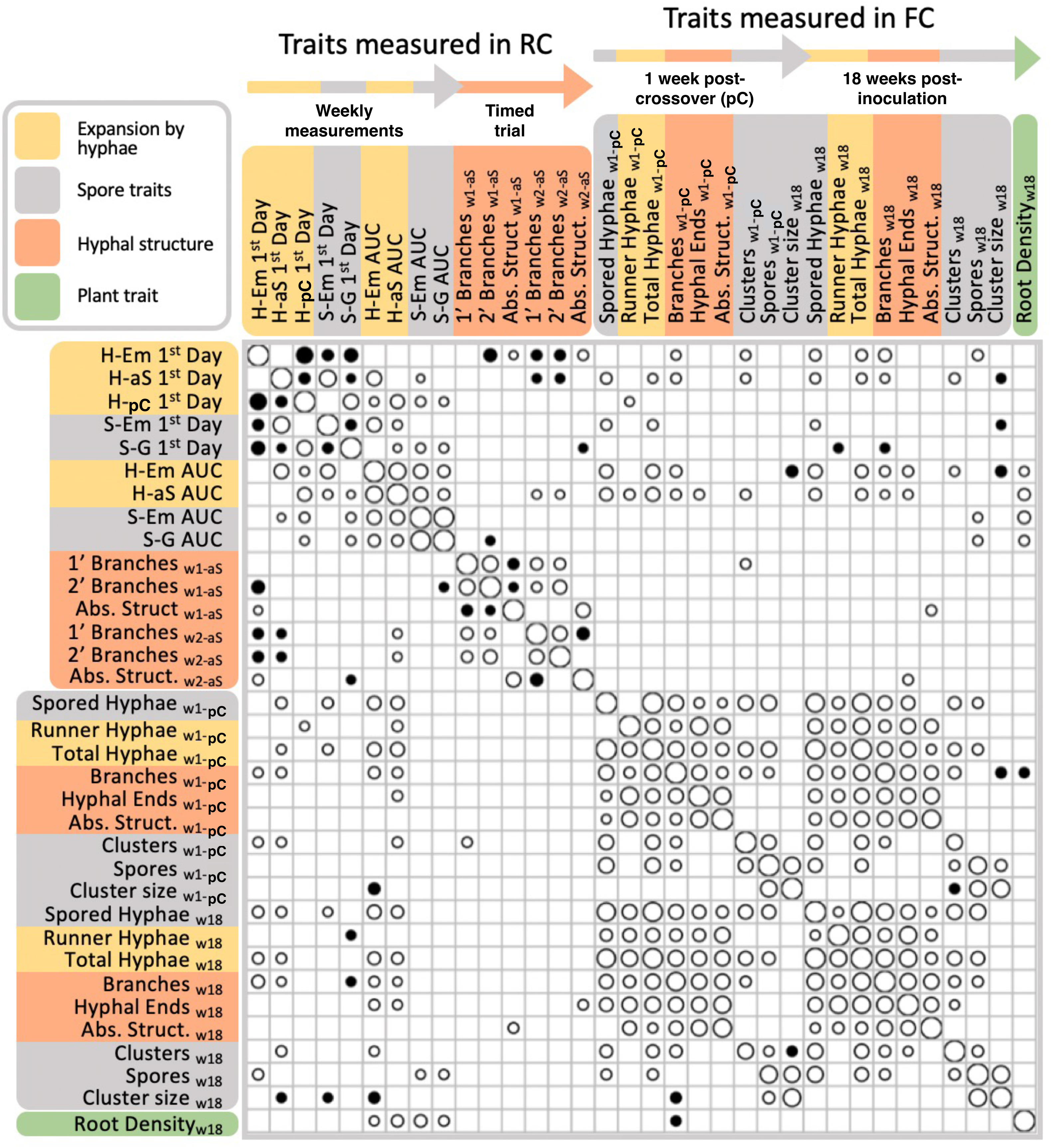
Correlogram of all measured traits, with the strength of the correlation visualized by the size of the circle (determined by *R*^2^; positive *R*^2^ white; negative *R*^2^ black). Correlations with *p* < 0.05 are not displayed. **RC**, root compartment; **FC**, fungal compartment; **H-Em**, number of emerging hyphae; **H-aS**, number of hyphae at the separator; **H-pC**, number of hyphae growing post-crossover, over the separator; **S-Em**, number of newly emerged spores; **S-G**, number of germinated spores; **1st Day**, first appearance; **AUC**, area under a growth curve; **w#-aS**, week number (one or two) of the first hypha touching the separator; **Abs. Struct.**, absorbance structures; **w1-pC**, one week post-crossover; **w18**, 18 weeks after RC inoculation.

**Figure 3.**
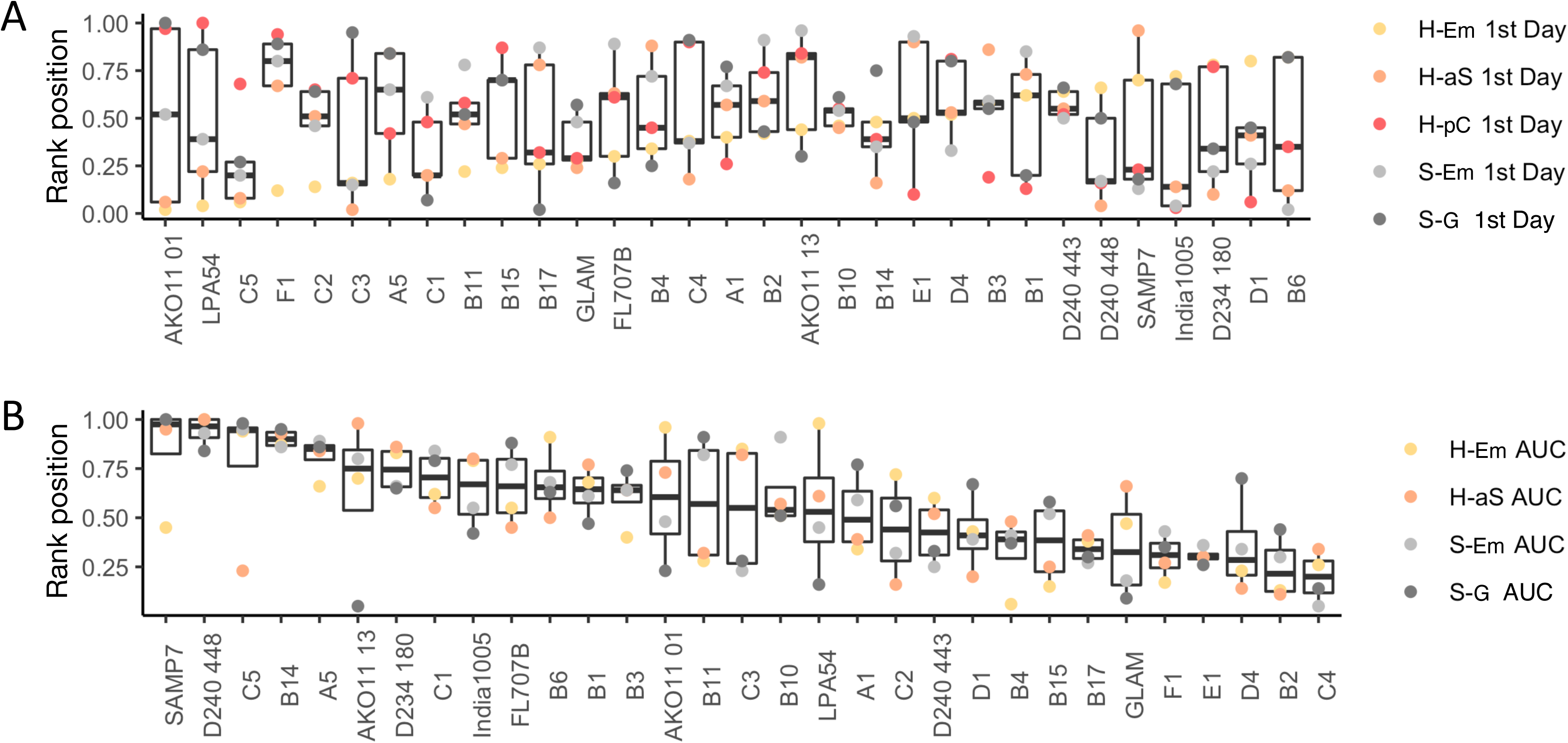
Profile of each isolate determined by rankings of a collection of RC traits. Low rankings (bottom of the y-axis, near 0) represent faster traits in **(A)** the 1^st^ day traits of hyphae emergence and extension, and in sporulation and germination. Lower ranks are faster growth rates for **(B)** area under the curve (AUC) quantifications. Isolates are ordered (x-axis) by those that were the fastest to slowest to emerge (**A**, left to right) and by their AUC trait median (**B**, left to right). Only isolates with at least two replicates remaining for each trait are displayed. Boxplots have median values indicated for all rankings in each trait group (**A**, 1^st^ day or **B**, AUC traits). **RC**, root compartment; **H-Em**, number of emerging hyphae; **H-aS**, number of hyphae at the separator; **H-pC**, number of hyphae post-crossover, growing over the separator; **S-Em**, number of newly emerged spores; **S-G**, number of germinated spores; **1st Day**, first appearance.

**Figure 4.**
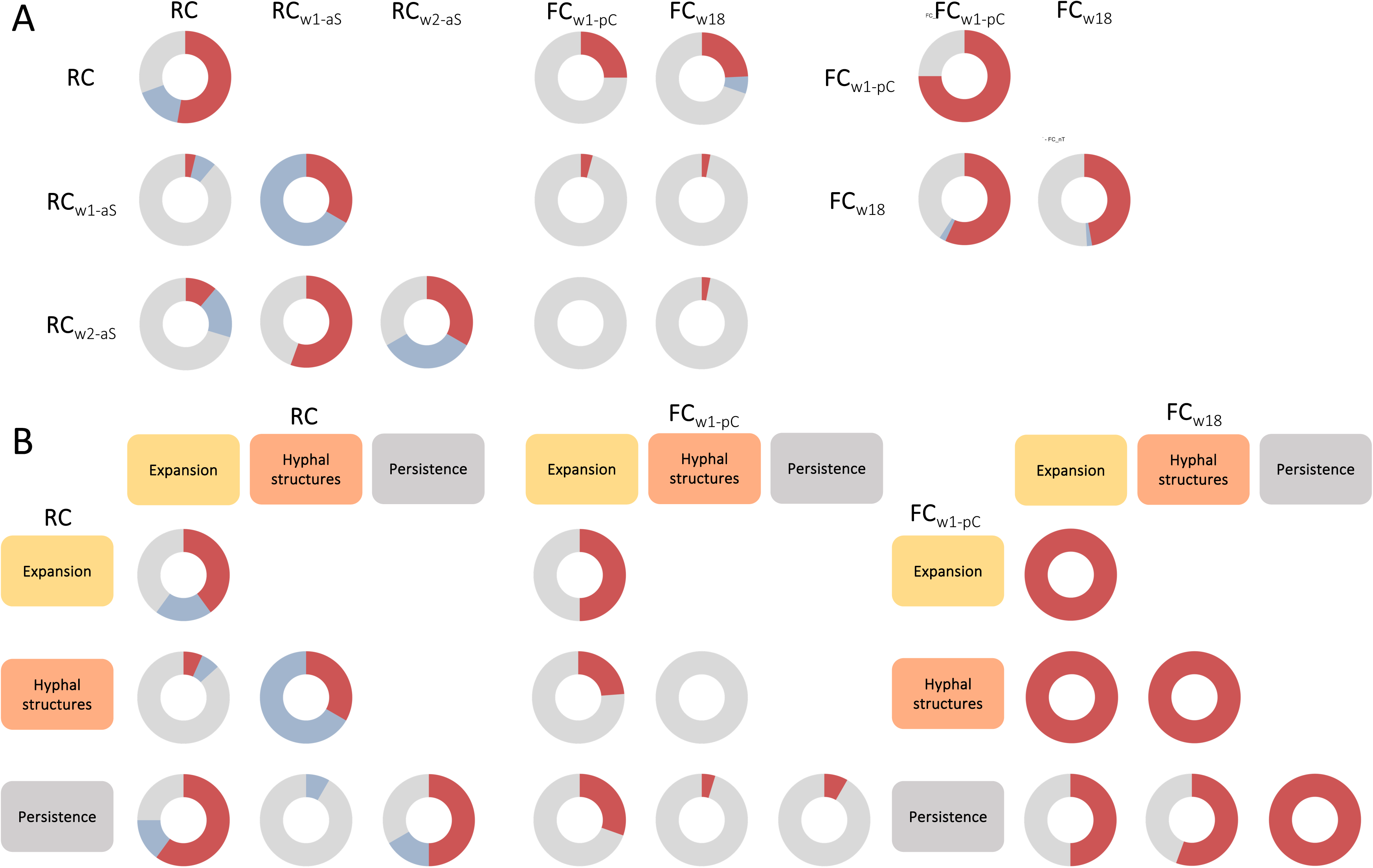
Summaries of the correlation results grouped by **(A)** experimental stage (non-timed trial measurements in the RC: **RC**; RC timed trial first timepoint: **RC**_w1-aS_; RC timed trial second timepoint: **RC**_w2-aS_; FC first timepoint, post-Crossover: **FC**_w1-pC_; FC second timepoint: **FC**_w18_), and **(B)** trait categories within experimental stage. In the latter, the RC timed trial is reduced to only the final measurements at the second timepoint (**RC**_w2-aS_ from (**A**)). Proportions of total correlations that were significantly positive (red), negative (blue), and non-significant (gray) are displayed in donut charts at the intersections of the various categories. **RC**, root compartment; **FC**, fungal compartment; **w#-aS**, week number (one or two) of the first hypha touching the separator; **w1-pC**, one-week post-Crossover, after the first hypha grew over the separator; **w18**, 18 weeks after RC inoculation.

**Figure 5.**
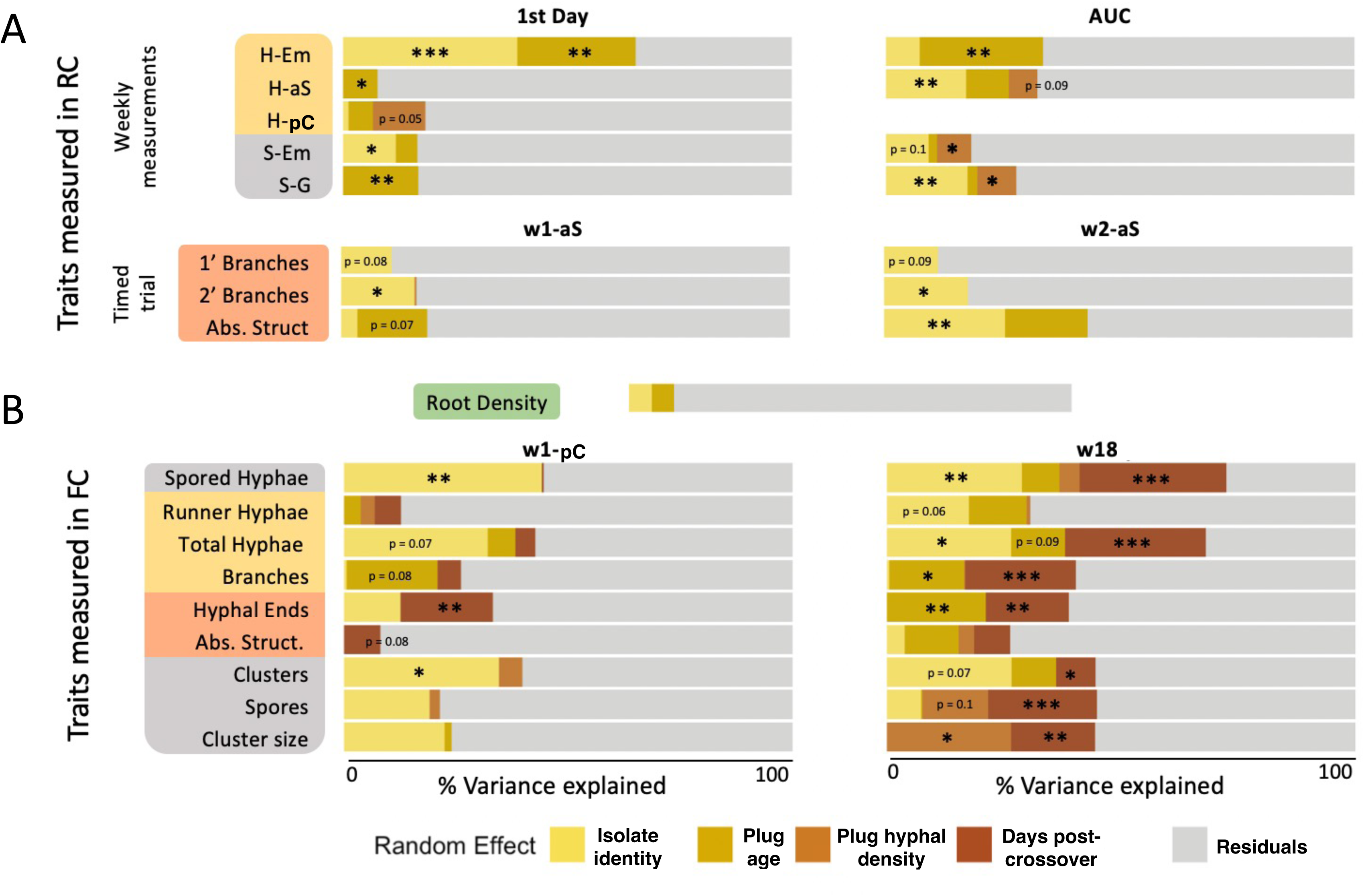
Variance decomposition (repeatability) results for **(A)** RC and **(B)** FC traits. Contributions of isolate identity (yellow), isolate plug age (gold), isolate plug hyphal density (light brown), and in (B), also day post-crossover per plate (dark brown) were analyzed for their contribution to trait variances. Residual variance is displayed in light gray. **RC**, root compartment; **FC**, fungal compartment; **1st Day**, first appearance; **AUC**, area under a growth curve; **w#-aS**, week number (one or two) of the first hypha touching the separator; **w1-pC**, one-week post-crossover, when the first hypha grew over the separator; **w18**, 18 weeks after RC inoculation. *** indicates *p* < 0.001; **, *p* < 0.01; *, *p* < 0.05; values with marginal significance are written in.

### Ranking of isolate traits for profile determination

To visualize trait diversity across isolates, all traits would need to be put into the same scale of interpretation (since they could be on quite different scales). A ranking system was established according to each isolate’s final average trait value (only for isolates with at least n = 2 plates remaining for all traits considered, Tab. S4). Trait ranks were between 0 (lowest, fastest trait values) to 1 (highest, slowest) and these were used for plotting (Fig. 3; medians of the ranks of all traits in a group were displayed).

### Statistical analysis

Variance decomposition was performed via the R package for repeatability, *rptR* [39] that estimates the proportion of total trait variance explained by random effects and their significance. We assessed contributions of random effects representing known groupings within the population of *R. irregularis* isolates (*e.g.*, isolate identity, initial plug differences in age, hyphal density) to the recorded fungal growth and reproductive traits (Fig. 5, Tab. S3, S5, S6). Remaining residual variance in each repeatability analysis represents unexplained variance among individuals.

## RESULTS

### Positive trait correlations dominate in R. irregularis, especially further from the plant

Significant correlations were more common among fungal compartment (FC) traits (99/153, 64.7%; bottom right of Fig. 2) compared to root compartment (RC) traits (46/105, 43.8%; top left of Fig. 2) and between-compartment traits (51/270, 18.8%). Negative correlations – indicating trade-offs – occurred mostly in RC correlations (14/46, 30.4%) and were rare for RC-FC (6/51, 11.8%) and FC-only (3/99, 3.0%) relationships. This suggests that *R. irregularis* trade-offs may primarily occur near roots.

The reduction of negative correlations in the FC does not appear to be due to the passage of time or changes in spatial constraints. To test this, we compared the two FC measurements: right after Crossover and at the end of the experiment. Among the two timepoints, a majority of the traits were correlated (53/64, 82.8%), and almost all positively (51/53, 96.2%). Within the first post-Crossover timepoint, 61.1% (22/36) of traits were correlated, all positively. For the end FC timepoint, all correlations (66.7%, 24/36) were also positive. These data indicate that FC traits relationships remain stable over time, and to saturation of the compartment.

### R. proliferus trait profiles show more opposite extremes than those of R. irregularis isolates

Trait profiles for each isolate were compared with the ranking analysis; several profiles were made for different collections of traits, such as the 1^st^ day group of traits (Fig. 3A). These profiles revealed whether isolates that quickly produced new hyphae (low rank for this trait) would always be fast in the appearance of all structures (consistently low rank). A larger interquartile range of the rankings would show that the traits of an isolate had various speeds relative to the population. *R. irregularis* isolates could also be compared to the two *R. proliferus* isolates with this analysis.

For 1^st^ day traits, it is clear that isolates that were first to produce new hyphae (low rank) did not have the same rank for all traits (Fig. 3A). The correlation analysis supports this, as other traits did not consistently correlate with fast emergence (Fig. 2); isolates could not be characterized as generally fast or slow. *R. irregularis* isolates exhibited large variation in 1^st^ appearance traits, as well as in rates of accumulation of hyphae and spores (AUCs, Fig. 3B), and FC traits (Fig. S2).

The two *R. proliferus* isolates (“AKO” isolates in Figs. 3, S2) often had some of the widest ranges in rankings within profile types and they were at the extremes of the rankings in comparison to the *R. irregularis* population. This may indicate that *R. proliferus* are subject to stronger trade-offs than isolates of *R. irregularis*, both in root– and non-root-influenced growing spaces.

### Early, near-root *R. irregularis* traits do not predict eventual trait values or relationships

*R. irregularis* hyphal and spore traits tend to correlate within the same compartment (RC or FC), but not between compartments (Fig. 4A, top row). It is to note that the main RC measurements (Fig. 2, traits without a subscript) only included hyphal expansion and spore traits, and the RC timed trial only included hyphal structure traits (w1-aS and w2-aS measurements in Fig. 2). This is in contrast to the two sets of FC measurements, which included all trait types in both sets (FC_w1-pC_ and FC_w18_). Therefore, the overview of RC to RC, RC to FC_w1-pC_, RC to FC_w18_ and among-FC correlations provided in the first row of Fig. 4A support only that there are fewer correlations among RC-FC hyphal expansion and spore traits, compared to within each compartment.

The summaries of significant relationships with RC_w1/2-aS_ traits shows that there are not many traits correlating with RC hyphal structure (primary, secondary, and high-density branching referred to as absorbance structures), including in the RC or to the FC, but hyphal structure traits do show strong internal trade-offs in both weeks (RC_w1-aS_-only and RC_w2-aS_-only; Fig. 4A). Supporting our previous findings (Fig. 2), the most overall correlations, and predominantly positive ones, are found among the FC traits (Fig. 4A, FC-only comparisons).

To investigate whether hyphal expansion and spore (persistence) traits may contain particular relationships within their trait categories, as seen for hyphal structural traits, we summarized correlations within all trait categories in and across each measurement periods (Fig. 4B). We found that most traits in the RC show correlations within their trait category. In addition, RC hyphal expansion and persistence traits also showed many cross-category correlations, which were mostly positive. Perhaps not surprisingly, hyphal structural traits did not often correlate across categories, even within the RC.

As the two within-trait-type FC measurement sets are also highly correlated (Fig. 4B, right), RC-FC correlations were only summarized for the RC and first FC dataset (Fig. 4B, middle). Most RC traits in all categories positively correlated with the early FC hyphal expansion traits. Persistence traits, like hyphal structure traits, did not correlate to the respective FC values. This suggests that RC traits can potentially predict hyphal expansion further from the host, but no other trait types. Though FC traits from the two measurements were highly correlated within type, persistence traits did have slightly less correlations, indicating that spore traits may be more influenced by time in the FC.

### Genetic individuality of isolates significantly explains variance in half of R. irregularis traits

While trait correlations help identify trade-offs, they do not reveal underlying influences on the traits. We performed a variance decomposition to determine the relative contributions of isolate identity (*i.e.*, genetics), and initial isolate subculture age and hyphal density, on RC traits from the *R. irregularis* population (Fig. 5A). For FC traits, we also tested days since Crossover (Fig. 5B).

How strongly a tested factor explains differences in trait values is represented by the strength of the significance. In the RC, isolate identity significantly explained variance in 10 of 15 traits, including 1^st^ day emergence traits but not 1^st^ day reaching the separator, of Crossover, or of spore germination. Of the four AUC traits, isolate identity significantly captured variance in number of hyphae at the separator, and numbers of new spores and spore geminations. Almost all hyphal structure traits of the timed trial were at least marginally significantly explained by isolate identity, which was interesting as these branching traits had the least connection to any other traits in our correlation analyses. Initial isolate subculture age influenced five traits, generally showing that older culture sources produced slower growth and reproductive production in the RC (Fig. S3A). Hyphal density in the original plug explained even fewer traits, where higher densities marginally linked to earlier presence at the separator, Crossover, and higher rates of spore appearance and germination (Fig. S3B).

In the FC, isolate identity significantly or marginally influenced 3 of 9 traits right after Crossover, including spored hyphae, total hyphae and spore clusters (Fig. 5B, left). Even though lacking in significance, large proportions of variance were attributed to isolate identity for hyphal ends, spores and cluster size. At the end of the experiment, isolate identity had a significant or marginally significant role in the same three traits, as well as for numbers of runner hyphae (4 of 9 traits; Fig. 5B, right). At this later time point, no additional traits had visual variance attributed to isolate identity. In general, branching traits in the FC were not explained by isolate identity, but were affected by the age of the isolate plug. Especially, older fungal plugs produced fewer total hyphae in the FC at the end of the experiment, with less branching but more frequent hyphal ends (Fig. S4A). There was no influence of the plug hyphal density on early FC traits, but higher density did significantly cause more numbers of spores and larger cluster sizes at the end of the experiment (Fig. S4B).

In the FC, days since Crossover was also tested and was a major factor in explaining 7 of 9 at the later timepoint (Fig. 5B, right). It additionally influenced two traits in the earlier FC timepoint: it explained variance in hyphal ends and absorbance structures (Fig. 5B, left). These traits appear highly sensitive to short-term changes, which at least for absorbance structures is consistent with the RC timed trial (Fig. 5A, bottom). Finally, we tested potential influences on the amount of carrot root present in the FC at the end of the experiment, but this was not explained by any factors tested (Fig. 5B, top). Overall, isolate identity, and thus genetic individuality, was the most consistent and significant driver of trait variation, especially in the RC and early FC. Later on in the FC, potentially confounded by day of Crossover, age and hyphal density had more effects.

## DISCUSSION

*In vitro* monitoring of *Rhizophagus irregularis* traits of a large population of isolates, grown individually across compartments with and without plant influence, revealed the variation, correlations among and consistency of these traits over time and in different environments (Figs. 1-4). This work addresses several decades of calls for a trait-based understanding of AMF growth from the fungal, rather than plant, perspective [11,23,38]. This joins recent works advancing our understanding of interspecific AMF spore traits [42,48] and intraspecific *R. irregularis* life history traits [36] and extends previous screens to allow a more detailed understanding of trade-offs in *R. irregularis* growth over time. Through a variance decomposition analysis, we additionally determined, for the first time, which *R. irregularis* traits are explained by genetic differences among isolates in comparison to other experimental influences. Isolate identity (*i.e.*, their genetics) was most influential in root compartment (RC) traits, where there were also more trait trade-offs.

*R. irregularis* is in the Glomeracae, characterized within Grime’s C-S-R framework as a ruderal (R) family known for high growth rates, early appearance of spores, short life spans, and dynamic hyphal structures with high recovery from disturbance [38]. Our work supports this: across the population of 48 *R. irregularis* isolates, there was generally an early appearance and germination of new spores (within 1-4 months) in comparison to what is known about other AMF species [43]. Spore appearance was weakly positively linked with growth rates (AUC of hyphal emergence and growth to the middle separator) in the RC, as well as with numbers of spored hyphae and total hyphae in the compartment without root influence (fungal compartment, FC; Fig. 2). We also found that if the starting fungal material for each isolate had larger amounts of subtending hyphae, there was faster growth and spore accumulation in the RC, and more spores in larger clusters in the FC (Fig. 5). This is expected for Glomeracae, since they are known for use of hyphal pieces to rapidly recover in disturbed environments.

*R. irregularis* isolates with high growth rates and early appearance of spores, however, were not always fast to begin their growth. The isolates demonstrate a large range of 1^st^ appearance of hyphae emerging from the initial fungal plug (1-4 months), unrelated to their eventual growth rates, and which was negatively correlated to 1^st^ appearance and germination of spores. This 1^st^ day hyphal trait had a stronger relationship with eventual hyphal branching traits, which is interesting as these traits (1’, 2’ branching, absorbance structures, hyphal ends) had little association with most other traits (Fig. 2, 4). Though age of the isolate material used for the experiment played a part in slowing the initial emergence and hyphal growth of isolates, these effects did not last for Crossover and beyond (Fig. 5). Our detailed phenotyping of 1^st^ day of appearance of traits and growth rates, and tracking these across temporal and environmental scales, thus allowed us to dissect relationships between influences on traits and the continuity of this influence throughout the isolate’s growth, as well as the consistency of trait relationships.

In the root-influenced RC, *R. irregularis* traits demonstrated the most trade-offs (negative correlations), though this represented only a third of significant relationships among traits in this compartment (Fig. 2). Most trait correlations, including the negative ones, were within trait types (Fig. 4). Previously, fungal traits have been separated into those for exploration (*e.g.*, expansion hyphae, without branching or spore clusters), those for local probing (structural hyphae, with various degrees of branching), and those for persistence (spore-related). Trait relationships that crossed trait categories included hyphal expansion traits correlating (mainly positively) with spore traits in the RC, but this changed to hyphal expansion with structure traits in the FC (Fig. 4). The two plate environments clearly create different trait relationships and growth patterns across *R. irregularis* isolates. This raises questions about what occurs in soil environments, where there could also be drastic gradients, possibly as abrupt as the differences in the plates, leading to different traits observed in the same isolate throughout time and space. Interestingly, the location in which hyphal structures were observed in the RC also influenced trait results (Tab. S5), supporting potential distance-related influences of plant roots as well as further AMF trait changes in a root-less environment.

From Crossover, nearly all trait correlations were positive in *R. irregularis* (Fig. 2). This did not seem to be the case for the comparison isolate species: *R. proliferus*, in which trade-offs among isolate traits in the FC were maintained (Fig. 3, Fig. S2). There were very minimal correlations among traits between the RC and FC (Fig. 4), with one notable exception: traits in all categories in the RC had some positive connection to the hyphal expansion traits in the FC (Fig. 4). It remains to be known if when a host plant experiences different stress factors, if the trends that we observed across the traits of the *R. irregularis* population, in the different environments, would remain consistent despite the host state. Additionally, as we saw that many traits are strongly influenced by the difference in isolates (extrapolated to their individual genetic make ups), it is interesting to determine in which traits a genome x environment interaction may play a role.

We provide, to our knowledge for the first time, information on the potential influence of individual isolate genetic identity on each trait, analyzed for traits in the RC, FC, and across time and space, in order to support this further research as well as to provide justification for potential forward genetic studies (like GWAS) to link traits to genetic drivers. Especially interesting for association mapping studies are the branching traits of *R. irregularis*, which although majorly unconnected to other traits, were explained strongly by isolate identity, and are essential in the interactions of this AMF with biotic and abiotic components in different soil locations. After this, mapping on hyphal expansion traits in spaces further from the roots, and numbers of spore clusters, could illuminate the genetics that determine the balance between exploration of *R. irregularis* and its production of its “insurance policy”, represented by spore structures.

Whether *R. irregularis* traits are able to be directly linked to gene candidates or not, their associations with strong correlation groups can also provide information on their potential for adaptability in a changing environment. Both genetic drivers and correlations of traits can inform adaptation: for example, highly correlated traits in plants have been shown to accelerate the adaptability of traits in changing environments if the correlation is in the direction of selection [44]. Positive correlations typically are more reinforcing to selection, though there can be exceptions depending on specificities of new environments. The prevalence of positive trait correlations found in our study and especially in the FC, further from the “controlled” space that the AMF shared with the root partner, is interesting in this context. Hyphal expansion traits had the most positive correlations, which may also indicate that they could adapt faster, in conjunction with strongly associated spore traits in the root-related space and with hyphal branching traits in the root-less space, which is where these latter traits did not have detectable influence from isolate identity. The absence of many negative trait correlations in this species suggests that there are few trade-offs that would likely slow down adaptability to a changing environment.

A putative mating-type (MAT) locus in *R. irregularis* was shown to have high diversity which matched genome divergence in this species [45]. It was interesting to investigate whether isolates with similar MAT types may group together in a non-metric multidimensional scaling (NMDS) analysis in terms of having a more similar trait profile than isolates of other MAT-types. Our analysis revealed that this was not the case (Fig. S1). In addition, it had been previously reported that homokaryotic and dikaryotic isolates of *R. irregularis* demonstrated different life history strategies [36]. We hoped to replicate this result by seeing a different trait space occupied by the different karyotype isolates in an NMDS, but they entirely overlapped (Fig. S1). We wondered if the fewer isolates used by Serghi *et al.* may have led to their conclusion, but when re-performing our analysis with only focus isolates including some homokaryons and dikaryons that they had used, we saw no significant differences between homokaryon and dikaryon trait profiles (Fig. S1). We therefore conclude that although our study supports that genetic differences among isolates influence at least half of the traits we measured in our experiment, they do not follow MAT-type or karyotype identifications.

All in all, we provide novel information in AMF trait-based research, including further support for and specifications on which traits are related to the genetic differences in isolates. We acknowledge that some AMF traits may manifest at the AMF community level, which was not within the scope of this study but is important to consider for future investigations. In addition, host plant state (*e.g.* stress and different growth responses) may alter the results we observed in this study, but the foundation is important for further investigations including biotic and abiotic stress factors from the plant and fungal sides. In our experiment, we did have the advantage of containing and standardizing the plant material, leading to a lack of difference in the final plant material per isolate plate that allowed us to decouple feedback effects in plant and AMF growth. Ideally, AMF growth traits such as the ones we measured would be observed in more natural conditions (soil set-ups). New technologies are allowing for visualization of *in vitro* fungal growth in a high-throughput manner both pre-symbiosis (microfluidic devices; [46]) and during symbiosis (automatic hyphal growth imaging and processing; [37]). In soil, combinations of new micromorphology techniques [47] and autofluorescence aspects of hyphae may be useful to track AMF in soil, but as of yet these would be destructive measurements. Regardless, there remains much work to do in defining AMF traits and life history and *in vivo* research with many isolates can push this dramatically forward.

## SUPPLEMENTARY INFORMATION

**Figure S1** NMDS analyses of trait spaces of isolates with different MAT– and karyo-types.

**Figure S2** FC traits ranking analyses after crossover (A, B) and experiment end (C, D).

**Figure S3** RC trait correlations with the (A) age and (B) hyphal density of the isolate plug.

**Figure S4** FC trait correlations with the (A) age and (B) hyphal density of the isolate plug.

**Table S1** Table of isolates used, in what rounds, and where they were previously characterized.

**Table S2** Metadata of the isolates used, including all potential random effects.

**Table S3** Repeatability analysis on select traits of isolates in both experimental rounds.

**Table S4** Isolate replication at key phenotyping timepoints.

**Table S5** Repeatability analyses on RC timed trial traits including the three view locations.

**Table S6** Repeatability analyses on FC traits including the different view locations.

## ACKNOWLEDGEMENTS

We thank the University of Lausanne, the organizational oversight of the Molecular Life Sciences Master’s program for support of J.V. and P.G. in this project, and the State of Vaud for their support of E.M.

## DATA AVAILABILITY

The raw data for this manuscript is available upon request.

## AUTHOR CONTRIBUTIONS

Conceptualization: J.V., E.M., I.S.; Methodology: J.V., E.M.; Formal Analysis: J.V., P.G., E.M.; Investigation: J.V., P.G.; Resources: I.S.; Data Curation: J.V.; Writing – Original Draft: J.V., E.M.; Writing – Review and Editing: J.V., P.G., E.M., I.S.; Visualization: J.V., E.M.; Supervision: E.M., I.S.; Project Administration: J.V., E.M., I.S.; Funding Acquisition: I.S.

## REFERENCES

1. Ma X, Xu X, Geng Q et al. Global arbuscular mycorrhizal fungal diversity and abundance decreases with soil available phosphorus. Global Ecol Biogeogr 2023;32:1423–34.

2. Smith SE, Smith FA. Roles of arbuscular mycorrhizas in plant nutrition and growth: new paradigms from cellular to ecosystem scales. Annu Rev Plant Biol 2011;62:227–50.

3. van der Heijden MGA, Martin FM, Selosse M et al. Mycorrhizal ecology and evolution: the past, the present, and the future. New Phytologist 2015;205:1406–23.

4. Yang G, Wagg C, Veresoglou SD et al. How soil biota drive ecosystem stability. Trends in Plant Science 2018;23:1057–67.

5. Jiang F, Zhang L, Zhou J et al. Arbuscular mycorrhizal fungi enhance mineralisation of organic phosphorus by carrying bacteria along their extraradical hyphae. New Phytologist 2021;230:304–15.

6. Kakouridis A, Hagen JA, Kan MP et al. Routes to roots: direct evidence of water transport by arbuscular mycorrhizal fungi to host plants. New Phytologist 2022;236:210–21.

7. Keymer A, Pimprikar P, Wewer V et al. Lipid transfer from plants to arbuscular mycorrhiza fungi. eLife 2017;6:e29107.

8. Bennett AE, Groten K. The costs and benefits of plant–arbuscular mycorrhizal fungal interactions. Annu Rev Plant Biol 2022;73:649–72.

9. Berger F, Gutjahr C. Factors affecting plant responsiveness to arbuscular mycorrhiza. Current Opinion in Plant Biology 2021;59:101994.

10. Lekberg Y, Jansa J, McLeod M et al. Carbon and phosphorus exchange rates in arbuscular mycorrhizas depend on environmental context and differ among co-occurring plants. New Phytologist 2024;242:1576–88.

11. Antunes PM, Stürmer SL, Bever JD et al. Enhancing consistency in arbuscular mycorrhizal trait-based research to improve predictions of function. Mycorrhiza 2025;35:14.

12. Genre A, Lanfranco L, Perotto S et al. Unique and common traits in mycorrhizal symbioses. Nat Rev Microbiol 2020;18:649–60.

13. Zhang S, Lehmann A, Zheng W et al. Arbuscular mycorrhizal fungi increase grain yields: a meta-analysis. New Phytologist 2019;222:543–55.

14. Qin M, Li L, Miranda J et al. Experimental duration determines the effect of arbuscular mycorrhizal fungi on plant biomass in pot experiments: a meta-analysis. Front Plant Sci 2022;13:1024874.

15. George NP, Ray JG. The inevitability of arbuscular mycorrhiza for sustainability in organic agriculture—a critical review. Front Sustain Food Syst 2023;7:1124688.

16. Lutz S, Bodenhausen N, Hess J et al. Soil microbiome indicators can predict crop growth response to large-scale inoculation with arbuscular mycorrhizal fungi. Nat Microbiol 2023;8:2277–89.

17. Ceballos I, Mateus ID, Peña R et al. Using variation in arbuscular mycorrhizal fungi to drive the productivity of the food security crop cassava. 2019, DOI: 10.1101/830547.

18. Ordoñez YM, Villard L, Ceballos I et al. Inoculation with highly-related mycorrhizal fungal siblings, and their interaction with plant genoptypes, strongly shapes tropical mycorrhizal fungal community structure. 2020, DOI: 10.1101/2020.07.31.230490.

19. Stahlhut KN, Conway M, Mason CM et al. Intraspecific variation in mycorrhizal response is much larger than ecological literature suggests. Ecology 2023;104:e4015.

20. Lee E-H, Eo J-K, Ka K-H et al. Diversity of arbuscular mycorrhizal fungi and their roles in ecosystems. Mycobiology 2013;41:121–5.

21. Sanders IR, Rodriguez A. Aligning molecular studies of mycorrhizal fungal diversity with ecologically important levels of diversity in ecosystems. The ISME Journal 2016;10:2780–6.

22. Marrassini V, Ercoli L, Kuramae EE et al. Arbuscular mycorrhizal fungi originated from soils with a fertility gradient highlight a strong intraspecies functional variability. Applied Soil Ecology 2024;197:105344.

23. Chaudhary VB, Holland EP, Charman-Anderson S et al. What are mycorrhizal traits? Trends in Ecology & Evolution 2022;37:573–81.

24. Aavik T, Träger S, Zobel M et al. The joint effect of host plant genetic diversity and arbuscular mycorrhizal fungal communities on restoration success. Functional Ecology 2021;35:2621–34.

25. Ossler JN, Heath KD. Shared genes but not shared genetic variation: legume colonization by two belowground symbionts. The American Naturalist 2018;191:395–406.

26. Koch AM, Kuhn G, Fontanillas P et al. High genetic variability and low local diversity in a population of arbuscular mycorrhizal fungi. Proc Natl Acad Sci USA 2004;101:2369–74.

27. Masclaux FG, Wyss T, Pagni M et al. Investigating unexplained genetic variation and its expression in the arbuscular mycorrhizal fungus *Rhizophagus irregularis*: a comparison of whole genome and RAD sequencing data. PLoS ONE 2019;14:e0226497.

28. Robbins C, Cruz Corella J, Aletti C et al. Generation of unequal nuclear genotype proportions in *Rhizophagus irregularis* progeny causes allelic imbalance in gene transcription. New Phytologist 2021;231:1984–2001.

29. Chaturvedi A, Cruz Corella J, Robbins C et al. The methylome of the model arbuscular mycorrhizal fungus, Rhizophagus irregularis, shares characteristics with early diverging fungi and Dikarya. Commun Biol 2021;4:901.

30. Dallaire A, Manley BF, Wilkens M et al. Transcriptional activity and epigenetic regulation of transposable elements in the symbiotic fungus *Rhizophagus irregularis*. Genome Res 2021;31:2290–302.

31. Yildirir G, Sperschneider J, Malar C M et al. Long reads and Hi-C sequencing illuminate the two-compartment genome of the model arbuscular mycorrhizal symbiont *Rhizophagus irregularis*. New Phytologist 2022;233:1097–107.

32. Davison J, Moora M, Öpik M et al. Global assessment of arbuscular mycorrhizal fungus diversity reveals very low endemism. Science 2015;349:970–3.

33. Öpik M, Moora M, Liira J et al. Composition of root-colonizing arbuscular mycorrhizal fungal communities in different ecosystems around the globe. Journal of Ecology 2006;94:778– 90.

34. Savary R, Masclaux FG, Wyss T et al. A population genomics approach shows widespread geographical distribution of cryptic genomic forms of the symbiotic fungus *Rhizophagus irregularis*. The ISME Journal 2018;12:17–30.

35. Kokkoris V, Miles T, Hart MM. The role of *in vitro* cultivation on asymbiotic trait variation in a single species of arbuscular mycorrhizal fungus. Fungal Biology 2019;123:307–17.

36. Serghi EU, Kokkoris V, Cornell C et al. Homo– and dikaryons of the arbuscular mycorrhizal fungus *Rhizophagus irregularis* differ in life history strategy. Front Plant Sci 2021;12:715377.

37. Oyarte Galvez L, Bisot C, Bourrianne P et al. A travelling-wave strategy for plant–fungal trade. Nature 2025;639:172–80.

38. Chagnon P-L, Bradley RL, Maherali H et al. A trait-based framework to understand life history of mycorrhizal fungi. Trends in Plant Science 2013;18:484–91.

39. Stoffel MA, Nakagawa S, Schielzeth H. rptR: repeatability estimation and variance decomposition by generalized linear mixed-effects models. Methods Ecol Evol 2017;8:1639–44.

40. Rosikiewicz P, Bonvin J, Sanders IR. Cost-efficient production of *in vitro Rhizophagus irregularis*. Mycorrhiza 2017;27:477–86.

41. Nakagawa S, Schielzeth H. Repeatability for Gaussian and non-Gaussian data: a practical guide for biologists. Biological Reviews 2010;85:935–56.

42. Chaudhary VB, Nokes LF, González JB et al. TraitAM, a global spore trait database for arbuscular mycorrhizal fungi. Sci Data 2025;12:588.

43. Oehl F, Sieverding E, Ineichen K et al. Distinct sporulation dynamics of arbuscular mycorrhizal fungal communities from different agroecosystems in long-term microcosms. *Agriculture*, Ecosystems & Environment 2009;134:257–68.

44. Etterson JR, Shaw RG. Constraint to adaptive evolution in response to global warming. Science 2001;294:151–4.

45. Lee S-J, Risse E, Mateus ID et al. Evolution of unexpected diversity in a putative mating type locus and its correlation with genome variability reveals likely asexuality in the model mycorrhizal fungus *Rhizophagus irregularis*. BMC Genomics 2024;25:888.

46. Richter F, Calonne-Salmon M, Van Der Heijden MGA et al. *AMF-SporeChip* provides new insights into arbuscular mycorrhizal fungal asymbiotic hyphal growth dynamics at the cellular level. Lab Chip 2024;24:1930–46.

47. Jiménez-Martínez A, Gutiérrez-Castorena MaDC, Montaño NM et al. Micromorphology and thematic micro-mapping reveal differences in the soil structuring traits of three arbuscular mycorrhizal fungi. Pedobiologia 2024;104:150953.

48. Aguilar-Trigueros CA, Hempel S, Powell JR et al. Bridging reproductive and microbial ecology: a case study in arbuscular mycorrhizal fungi. The ISME Journal 2018;13;873–84.

