## Supplementary Information for "Dissecting mycorrhizal fungal trait variation, its genetic basis and trade-offs"

The following Supplementary Information is available for this article:

**Figure S1** NMDS analyses of trait spaces of isolates with different MAT- and karyo-types.

**Figure S2** FC traits ranking analyses after crossover (A, B) and experiment end (C, D).

**Figure S3** RC trait correlations with the (A) age and (B) hyphal density of the isolate plug.

**Figure S4** FC trait correlations with the (A) age and (B) hyphal density of the isolate plug.

**Table S1** Table of isolates used, in what rounds, and where they were previously characterized.

**Table S2** Metadata of the isolates used, including all potential random effects.

**Table S3** Repeatability analysis on select traits of isolates in both experimental rounds.

**Table S4** Isolate replication at key phenotyping timepoints.

**Table S5** Repeatability analyses on RC timed trial traits including the three view locations.

**Table S6** Repeatability analyses on FC traits including the different view locations.

**Figure S1** NMDS analyses of the isolates grouped by MAT- (top panels) and karyo-type (middle and bottom panels) using **(A)** root compartment (RC) and **(B)** fungal compartment (FC) traits. The FC traits were from the time-adjusted measurement, 1 week after Crossover. The middle panels show the karyotype comparison by all homokaryon and dikaryon lines with complete trait profiles. The lower panels are with two homokaryon and two dikaryon lines for an even comparison. Each homokaryon shares a MAT-type with one of the dikaryons (Tab. S2). *n.s.* = non-significant distinction of plate trait profiles by an ANOSIM test in R (*vegan* package).

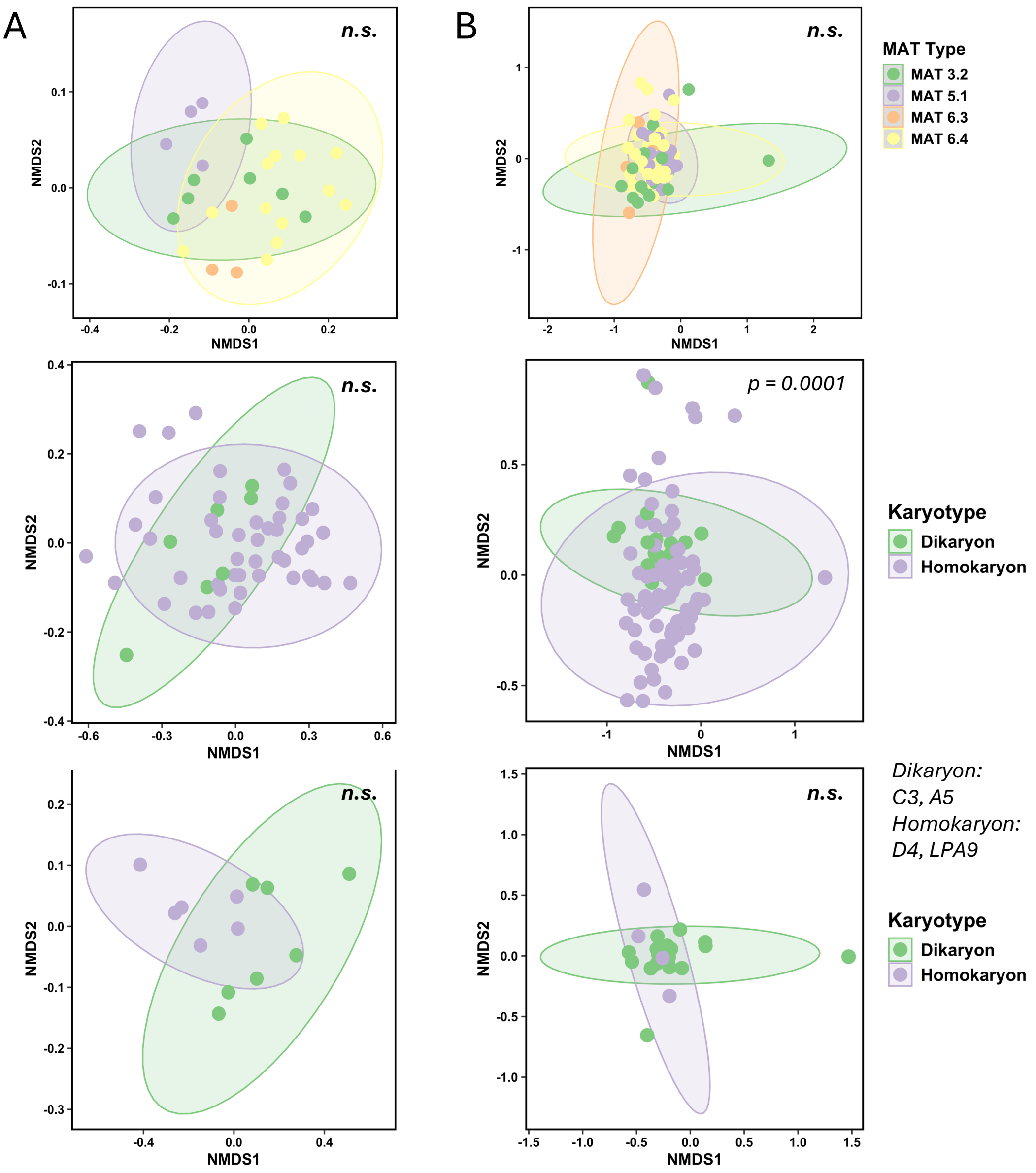

**Figure S2** FC traits ranking analyses 1 week after Crossover (A, B) and at the 18-week post-inoculation timepoint, without timing adjustment, at the experiment end (C, D). Lines plotted had more than 2 replicates remaining at the respective phenotyping timepoint (Tab. S4).

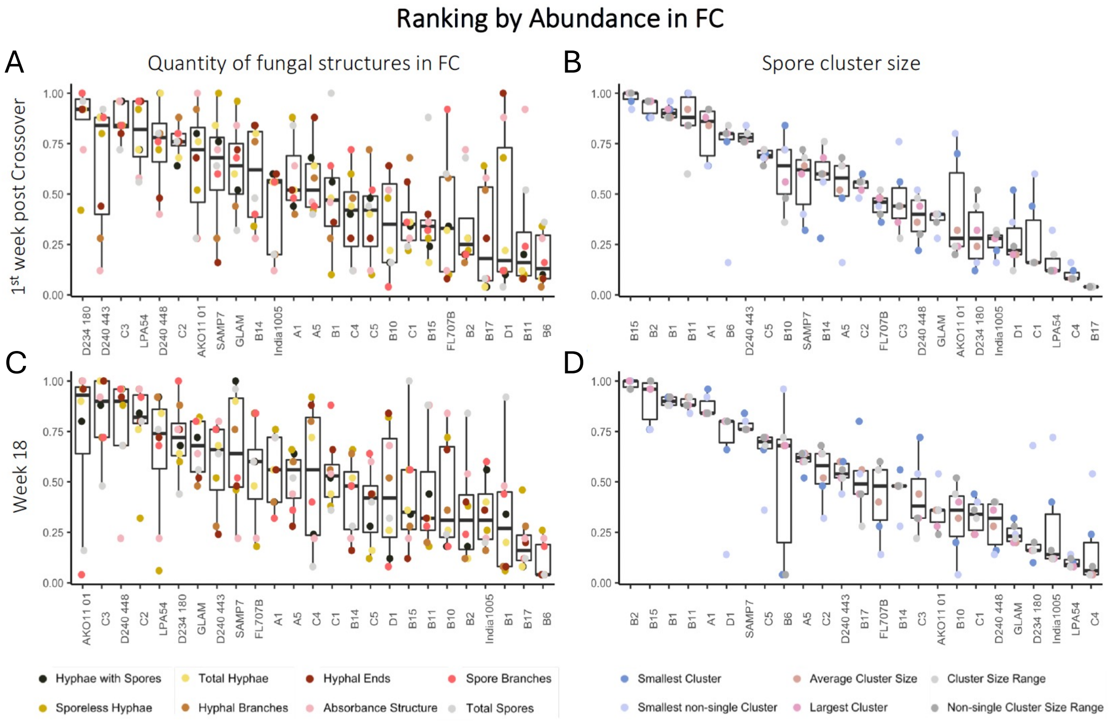

**Figure S3** RC trait correlations (Pearson's) with the (A) age and (B) hyphal density of the original isolate plug. Only traits that had significant contributions from age or hyphal density in the variance decomposition analyses (Fig. 5) are included. R and *p* values are displayed.

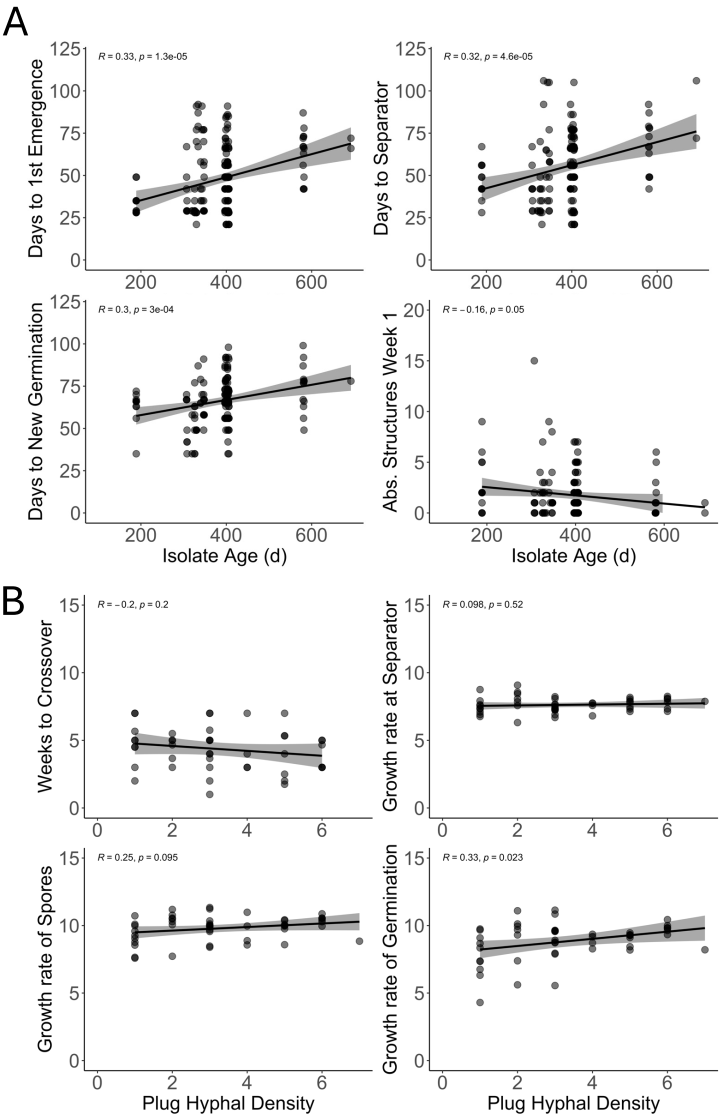

**Figure S4** FC trait correlations (Pearson's) with the (A) age and (B) hyphal density of the original isolate plug. Only traits that had significant contributions from age or hyphal density in the variance decomposition analyses (Fig. 5) are included. R and *p* values are displayed.

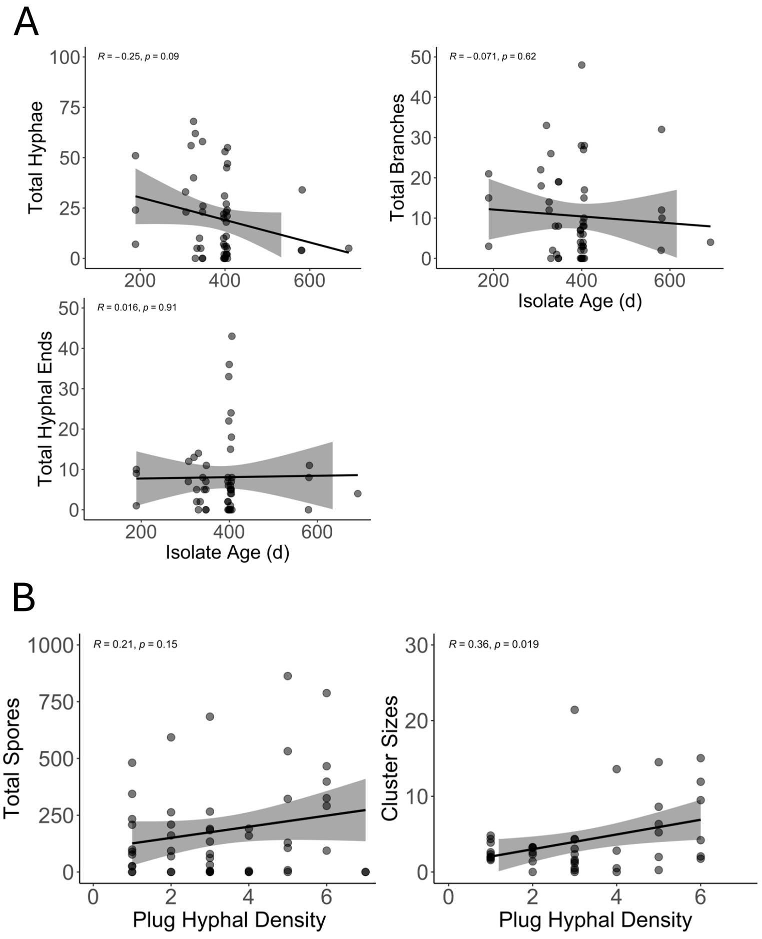

**Table S1** Table of isolates used, in what rounds, and where they were previously characterized. If the source for the previous characterization is blank, it is the same as the last source named above.

***Rhizophagus irregularis***

| Isolate name | Round 1 | Round 2 | Previous characterization |
|---|---|---|---|
| A1 | X | X | Savary et al. 2018 – *The ISME Journal* |
| A2 | X | | |
| A3 | X | | |
| A5 | X | | |
| B1 | X | | |
| B2 | X | | |
| B3 | X | | |
| B4 | X | | |
| B6 | X | | None, characterized in this study |
| B7 | X | | Savary et al. 2018 – *The ISME Journal* |
| B8 | X | X | |
| B10 | X | | |
| B11 | X | | |
| B12 | X | | |
| B14 | X | | |
| B15 | X | | |
| B17 | X | | |
| BEG53 (LPA30) | X | | |
| C1 | X | | |
| C2 | X | X | |

| | | | |
|---|---|---|---|
| C3 | X | | |
| C4 | X | | |
| C5 | X | | |
| CAN (DAOM197198-CH) | X | X | |
| D1 | X | X | |
| D2 | X | | |
| D3 | X | | |
| D4 | X | | |
| DAOM234180 | X | X | |
| DAOM234328 | X | | |
| DAOM240158 | X | | |
| DAOM240409 | X | | |
| DAOM240443 | X | | |
| DAOM240448 | X | | |
| DAOM240721 | X | X | |
| DAOM Miroslav (DAOM197198-CZ) | X | X | |
| DHP13 | X | | Rincón et al. 2021 - *Mycorrhiza* |
| E1 | X | | Savary et al. 2018 – *The ISME Journal* |
| ESLQ569 (ESLQ69) | X | | |
| F1 | X | | |
| FL707B (FL707) | X | | |
| G1 | X | | |
| GLAM | X | X | |
| India1005 | X | | Lee et al. 2024 – *BMC Genomics* |

| | | | |
|---|---|---|---|
| KUVA | X | | Savary et al. 2018 – *The ISME Journal* |
| LPA9 | X | | |
| LPA54 | X | | |
| SAMP7 | | X | |

***Rhizophagus proliferus***

| **Isolate name** | **Round 1** | **Round 2** | **Previous characterization** |
|---|---|---|---|
| AKO11 01 | X | | Savary et al. 2018 – *The ISME Journal* |
| AKO11 13 | X | | |

**Table S2** Metadata of the isolates used including all potential random effects.

*Submitted as an Excel file.*

**Table S3** Repeatability analysis on select traits of isolates in both experimental rounds.

*Submitted as an Excel file.*

**Table S4** Isolate replication at key phenotyping timepoints.

*Submitted as an Excel file.*

**Table S5** Repeatability analyses on RC timed trial traits including the three view locations.

*Submitted as an Excel file.*

**Table S6** Repeatability analyses on FC traits including the different view locations.

*Submitted as an Excel file.*

### References


Savary R, Masclaux FG, Wyss T *et al.* A population genomics approach shows widespread geographical distribution of cryptic genomic forms of the symbiotic fungus *Rhizophagus irregularis*. *The ISME Journal* 2018;12:17–30.

Rincón C, Droh G, Villard, L *et al.* Hierarchical spatial sampling reveals factors influencing arbuscular mycorrhizal fungus diversity in Côte d'Ivoire cocoa plantations. *Mycorrhiza* 2021;**31**:289.

Lee S-J, Risse E, Mateus ID *et al.* Evolution of unexpected diversity in a putative mating type locus and its correlation with genome variability reveals likely asexuality in the model mycorrhizal fungus *Rhizophagus irregularis*. BMC Genomics 2024;25:888.
